# Quantum Deep Learning Pipeline for Next Generation Network Biology

**DOI:** 10.1101/2025.10.28.685074

**Authors:** Milad Yazdi, Kushala Ralpati Srinivas, Ajay Yadav

## Abstract

**Purpose:** Module discovery in omics networks is central to interpretation. Classical pipelines capture broad community structure, but exact search for small, connected, topology aware modules is combinatorial, making exhaustive solutions impractical at genome scale. Moreover, most quantum clique formulations to date optimize only maximal density on non biological graphs, which mismatches the heterogeneous shapes of real biological modules.

**Methods:** We trained a symmetric 10-dimensional autoencoder on GTEx normals (Heart Left Ventricle; Muscle Skeletal; UCSC Xena Toil) to obtain tissue-specific latent representations. For each tissue we built a 10*10 latent-node correlation graph and formulated a QUBO that rewards strong edges and penalizes isolated selections (edge threshold τ=0.80). QAOA (depth=3, COBYLA) generated high-probability bitstrings, which we post-filtered to retain connected, non-overlapping subsets; decoder triggered gene sets and multi-library enrichment provided functional interpretation.

**Results:** QAOA distilled the Heart graph to three dyads (2-node modules with strong-edge sums ≈0.89/0.95), pairing conduction with mitochondria, conduction with ECM/adhesion, and two contractile nodes. In Muscle, QAOA returned one dominant 6-node module (edge-sum ≈7.0) integrating contractile machinery, electrophysiology, mitochondrial metabolism, and translation; a weaker ECM leaning triplet was visible but fell below the top 10 threshold.

**Conclusion:** QAOA based quantum optimization yields discrete, testable modules from latent correlations that match tissue programs. Despite a 10-node demo (current limits), the scale-agnostic design extends to 10^3 10^5 nodes and multi omics via hierarchical compression and hybrid search supporting quantum modeling for next generation network biology.

## Introduction

Biological systems are often represented as networks in which genes, proteins, and metabolites are nodes connected by edges denoting co-expression, regulation, or physical interaction. These networks exhibit a modular organization: densely connected groups of nodes correspond to functional units such as pathways, complexes, or regulatory circuits [1]. Detecting such modules is central to systems biology, as it enables interpretation of high-dimensional omics data in terms of cellular function and disease mechanisms [2]. Classical approaches such as WGCNA, community detection with Louvain or Leiden algorithms [3, 4], and graph-based searches like clique enumeration or Steiner trees have been widely applied. However, these methods face intrinsic limitations. Optimizing modularity is NP-hard, leading to reliance on heuristics [5]. Greedy clustering struggles with the “resolution limit,” where smaller but biologically relevant modules are missed in large networks. Likewise, finding maximum-weight or dense subgraphs is equivalent to NP-complete problems, and becomes intractable at genome scale. With modern datasets now spanning tens of thousands of genes or single-cell profiles, scaling module discovery remains a fundamental challenge.

Quantum computing has emerged as a candidate to address such combinatorial bottlenecks. Many module-finding tasks can be formulated as quadratic unconstrained binary optimization (QUBO), where binary variables encode gene inclusion and the objective rewards intra-module connectivity while penalising isolation [6]. This formulation maps directly to the Ising model and can be solved by the Quantum Approximate Optimization Algorithm (QAOA) [7]. QAOA is a hybrid variational algorithm in which parameterised quantum circuits approximate the ground state of the problem Hamiltonian, yielding high-quality solutions to NP-hard problems with relatively shallow circuits [8]. Although QAOA has shown promise on benchmark problems such as Max-Cut and scheduling, applications to biological networks are almost entirely unexplored. Current implementations remain constrained by device noise and limited depth, but advances such as warm-start QAOA and recursive decomposition suggest practical scaling pathways [9]. The gap between potential and application highlights both the opportunity and the need to test QAOA within realistic network biology pipelines.

Recent work has begun to operationalise clique- and subnetwork-centric objectives on quantum or quantum-inspired hardware. Photonic/analog XY-Ising machines (“DOMINO”) have been used to extract maximum-weighted gene cliques from large co-expression graphs (e.g., a 7-gene regulator set from a 386-node K562 network;[10]. In genetics, epistasis detection has been cast as a clique-style QUBO with a network-medicine prior, enabling hybrid/annealing solvers to target disease-associated SNP subnetworks [11]. Gate-model progress includes QOMIC, a quantum optimization approach that identifies recurrent network motifs in human GRNs across neurodegenerative diseases [12]. Relatedly, hybrid annealing has been applied to residue-interaction networks to pinpoint central “hub” residues (RinQ)[13]. At the algorithmic level, Seidel-matrix continuous-time quantum walks have been proposed as a parameter-free route to maximum clique [14]. Together these studies reinforce the viability of QUBO/Ising mappings for biological graphs and motivate our QAOA-based search for dense, connected latent-space modules as a complementary, data-driven testbed.

Here we present a proof-of-concept workflow that couples deep representation learning with quantum optimisation. We trained a nonlinear autoencoder on transcriptomic data from Heart (Left Ventricle) and Muscle (Skeletal), producing a 10-dimensional latent representation. Tissue-specific latent correlation graphs (10 * 10) were then constructed and encoded as QUBO problems, with objectives rewarding strong edges and penalising isolated nodes. QAOA was applied with stringent post-filters to retain only connected, non-overlapping subgraphs. In Heart, this procedure yielded three dyadic modules, while in Muscle it recovered a single 6-node clique integrating contractile, electrophysiological, mitochondrial, and translational functions. To our knowledge, this is among the first demonstrations of QAOA applied to a learned biological latent graph. Although our current study is restricted to 10 nodes, the approach is scale-agnostic: latent compression, block decomposition, and hybrid classical–quantum strategies could extend the method to thousands of nodes and to multi-omics integration. By linking representation learning with quantum optimisation, we establish a foundation for quantum-assisted module discovery in next-generation network biology.

## Materials and Methods

### Data sources and phenotype selection

We used the UCSC Xena Toil compendium to obtain the unified TPM expression matrix (TcgaTargetGtex_rsem_gene_tpm) and the companion phenotype table (TcgaTargetGTEX_phenotype.txt). From the phenotype, we explicitly selected normal GTEx samples for two tissues Heart – Left Ventricle and Muscle – Skeletal by robust string-matching across multiple candidate columns, and then balanced the cohorts by truncating to the same number of samples per tissue. To ensure downstream reproducibility, the final sample IDs used for each tissue were written to samples_Heart_LeftVentricle.csv and samples_Muscle_Skeletal.csv.

### Expression preprocessing

From the expression matrix we retained only the selected samples, filtered out zero-sum genes, and kept at most the top 5,000 most variable genes (otherwise all genes if fewer than 5,000). We then z-scored each gene across samples using StandardScaler (mean 0, variance 1). These steps yielded two aligned matrices expr_A (Heart) and expr_B (Muscle) for model training and analysis.

### Deep autoencoder: architecture, training, and exports

We trained a symmetric, fully connected 10-dimensional autoencoder on the z-scored expression (all selected samples combined). The encoder comprised Dense layers of sizes 2048, 1024, and 512 with ReLU activations, Dropout (0.2), and a BatchNorm before the linear latent layer (size = 10). The decoder mirrored the encoder (512→1024→2048→output). The network was optimized with Adam on MSE reconstruction loss (batch 32, up to 120 epochs) with early stopping (patience 12) using a 90/10 train/validation split; training curves (including a held-out test MSE line) were saved as fig_ae_training_curve.png. After training, we saved the full model (autoencoder_model.h5) and exported the encoder and decoder as separate artifacts (encoder.h5, decoder.h5) for reuse.

### Latent embeddings and PCA

We projected all samples to the 10-D latent space using the trained encoder and saved the table (samples ^*^10 nodes) as latent_nodes_^*^.csv per tissue. For visualization we performed PCA to 3 components on the latent matrix and saved pca3_^*^.csv and a 2-D scatter (fig_latent_pca.png).

### Tissue-specific 10^*^10 correlation matrices

For each tissue (Heart, Muscle) we computed the Pearson correlation matrix across the 10 latent nodes (size 10^*^10), clipped to [−1, 1], saved the matrices as corr10×10_heart_leftventricle.csv and corr10×10_muscle_skeletal.csv, and exported annotated heatmaps (fig_corr_^*^.png). These matrices constitute the weighted graphs used in the quantum optimization stage.

### Decoder-triggered “core” gene sets and enrichment

To probe what each latent node encodes, we generated a baseline decoder output at the all-zeros latent vector, then independently “activated” each latent dimension (+1.0) and recorded the absolute change in reconstructed expression relative to baseline. For each node we defined the core gene set as the union of the top-250 genes by absolute change and the top 5% by percentile threshold; we exported Ensembl IDs and mapped gene symbols with MyGene (node2genes_core/^*^_core_ids.csv, ^*^_core_symbols.csv). We performed multi-library enrichment with Enrichr (KEGG 2021, Reactome 2022, GO BP/MF/CC 2021, DisGeNET, ChEA 2022, DSigDB) and saved per-node CSVs.

### Quantum optimization of dense subgraphs (QAOA on QUBO)

#### Graph construction and normalization

For each tissue, we converted the 10^*^10 correlation matrix to a non-negative weight matrix by taking absolute values and normalizing to [0, 1] (normalize_matrix). A fixed edge threshold τ=0.80 determined “strong” edges.

#### QUBO formulation

We formulated the subset-selection objective as a QUBO minimization over binary variables xi∈{0,1}x_i\in\{0,1\}xi∈{0,1} (select node i) with three terms:

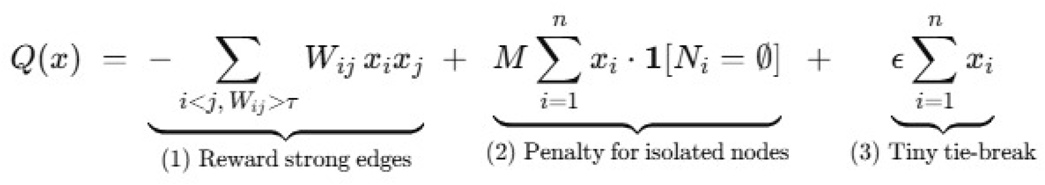

where the first term rewards strong edges among selected nodes, the second imposes a penalty M=10 on selecting any node whose set of strong neighbors NiN_iNi is empty (i.e., globally isolated at threshold τ), and □=10−3\epsilon=10^{-3}□=10−3 provides a tiny tie-break on set size. This QUBO was assembled using QuadraticProgram, then converted to an Ising operator for quantum processing.

#### QAOA solver and post-filtering

We ran QAOA at depth reps=3 with COBYLA (maxiter=80) as the classical optimizer, using the Aer Sampler with 10,000 shots and a fixed seed (42). From the measured bitstring distribution we retained up to 200 high-probability candidates. To reduce the search space by two orders of magnitude (from 1024 possible 10-node subsets to only 10 retained), we focused on the top-10 candidates by score for downstream analysis. Post-filters were applied in two stages: (i) candidate subsets had to contain ≥2 nodes and form a single connected component with respect to edges Wij>τW_{ij}>\tauWij>τ; (ii) a deduplication pass ensured that each node appeared in at most one accepted subset, retaining its highest-scoring occurrence. The final score for each subset was defined as the sum of strong-edge weights inside the subset. The resulting outputs were written to qaoa_top_heart_connected.csv and qaoa_top_muscle_connected.csv, alongside bar-plot summaries.

#### Software environment and versions

The autoencoder workflow depends on pandas≥2, numpy≥1.26, scikit-learn≥1.3, tensorflow 2.x, matplotlib, seaborn, gseapy, and mygene; scripts download inputs automatically and are Colab-ready. The quantum stage uses qiskit, qiskit-aer, qiskit-algorithms, and qiskit-optimization; a pinned configuration (qiskit==1.2.4, qiskit-aer==0.14.2, qiskit-algorithms==0.3.0, qiskit-optimization==0.6.1) is provided to ensure compatibility.

#### Reproducibility

All randomization steps were fixed by seeds (numpy/TensorFlow seeds = 0 for autoencoder; Qiskit global seed = 42) and all intermediate artifacts (sample lists, models, latent tables, PCA tables, correlation matrices, QAOA outputs) are written to disk for auditability and reuse.

## Results

### 3.1 Autoencoder training performance

The 10-dimensional autoencoder converged stably within ∼30 epochs. Training and validation MSE dropped steeply during the first ∼10 epochs and then plateaued with only minor oscillations (Fig. 1). The held-out test error (dashed line) was MSE = 0.348, closely matching the late-epoch validation range and indicating no material overfitting. The final generalization gap between training and validation remained modest, consistent with a well-fit model that captures shared structure without memorization. Overall, these diagnostics support the suitability of the learned latent space for downstream analyses.

**Figure 1:**
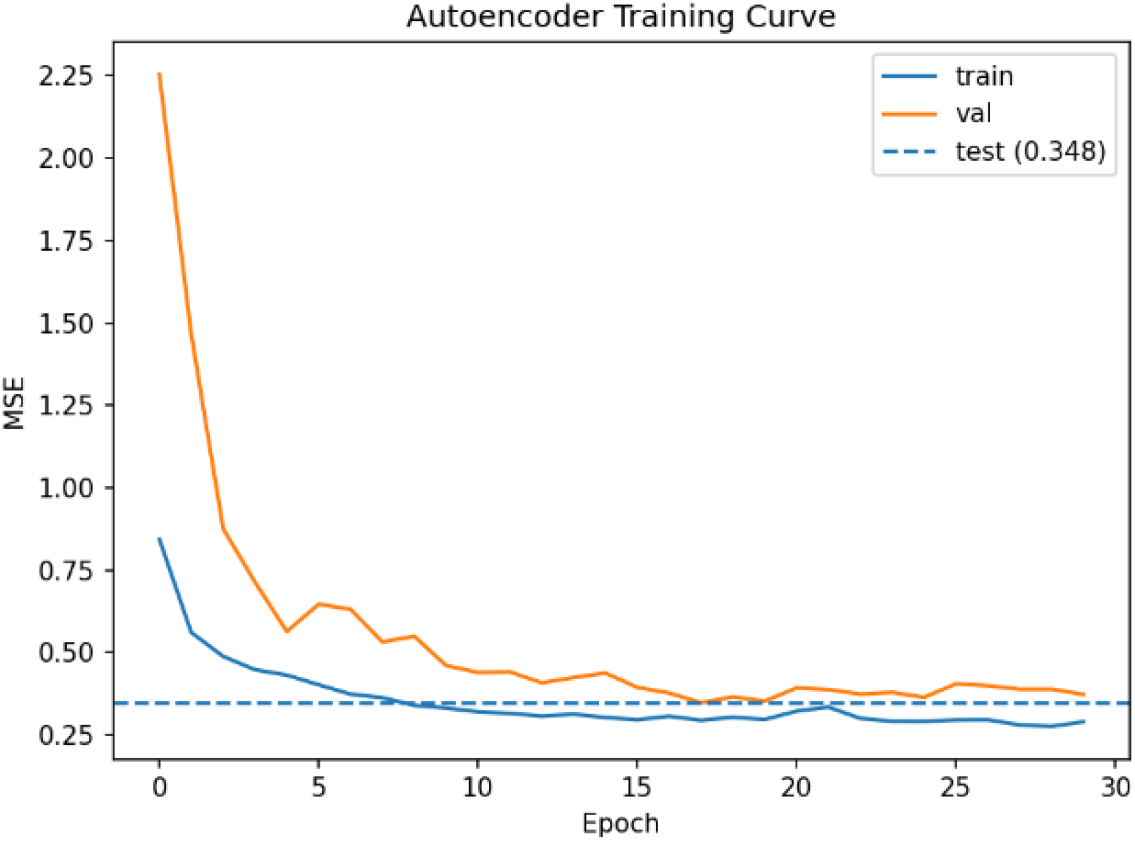
Training curve of the 10-dimensional autoencoder. Mean squared reconstruction error (MSE) is shown across epochs for training (blue) and validation (orange) sets, with the dashed line indicating the held-out test error (MSE = 0.348). Both curves converge smoothly, demonstrating stable training and good generalization.

Projecting all samples into the 10-dimensional latent space yielded a clear tissue-level structure. A 2-D PCA of the latent embeddings showed a pronounced separation of the two balanced cohorts primarily along PC1, with minimal overlap between groups (Fig. 2). The Heart cohort formed a compact cluster with a few outliers, whereas the Muscle cohort displayed broader dispersion along PC2, indicating greater within-group heterogeneity. Because labels were not used during training, this separation suggests that the autoencoder captured robust, tissue-specific variance suitable for downstream analyses.

**Figure 2:**
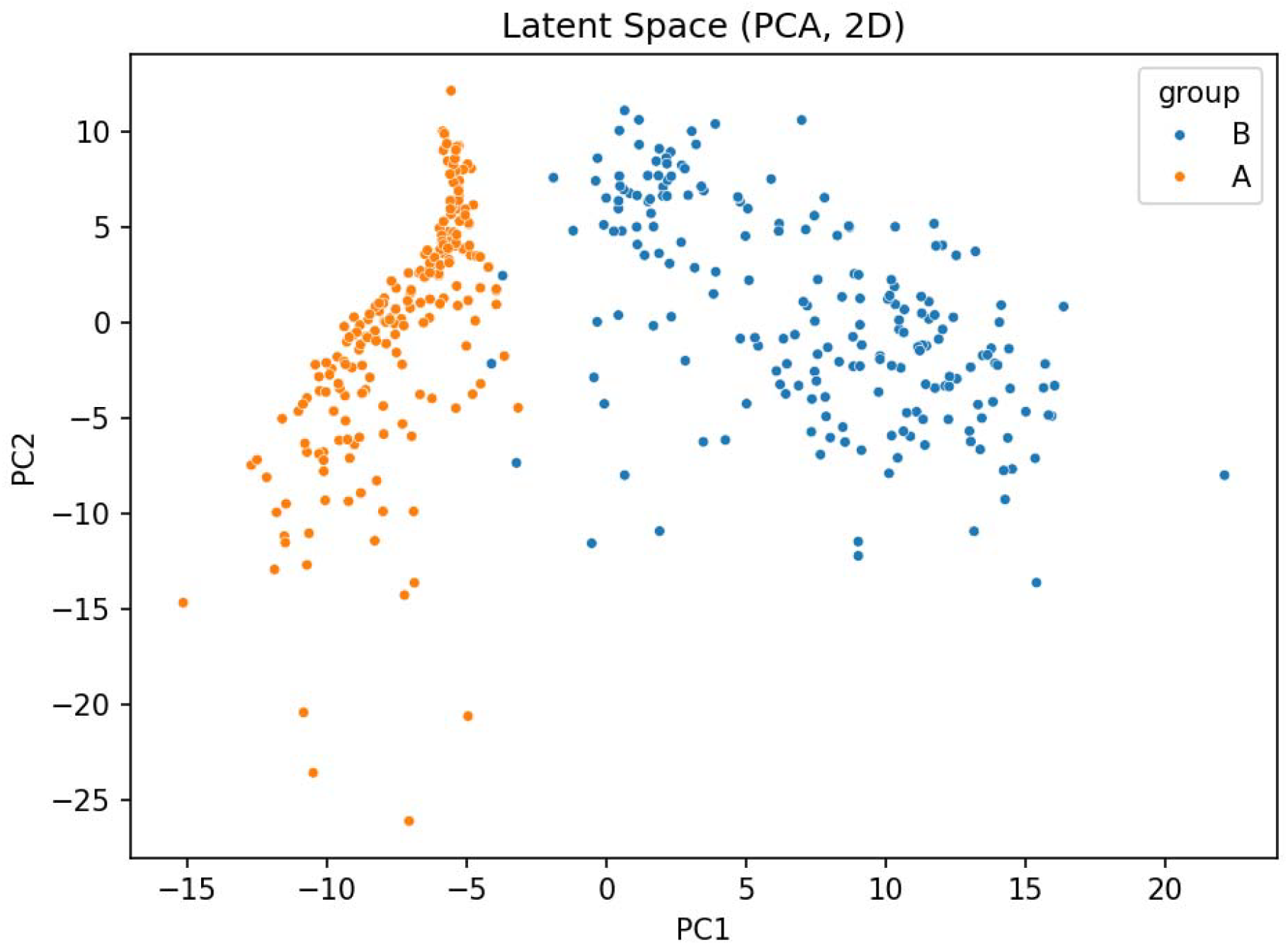
Latent space representation of Heart (Group A, orange) and Muscle (Group B, blue) samples projected into two principal components. The cohorts separate primarily along PC1, with Heart forming a compact cluster and Muscle showing broader dispersion, indicating robust tissue-specific structure captured by the autoencoder.

To summarize interactions among latent factors, we computed tissue-specific Pearson correlation matrices (10 10). The Heart matrix featured a mixture of strong positive and negative couplings, including node3–node9 (≈0.91) and node6–node7 (≈0.80) versus marked anti-correlations such as node4–node8 (≈ −0.95) and node1–node7 (≈ −0.89) (Fig. 3A). The Muscle matrix exhibited more numerous high-magnitude associations, for example node8–node9 (≈0.91), node6–node10 (≈0.90), node1–node8 (≈0.88), and strong antagonism between node3–node6 (≈−0.96) and node7 with nodes 3 and 9 (≈−0.90 and ≈−0.82) (Fig. 3B). These tissue-specific correlation patterns define the weighted graphs used in the subsequent optimization stage.

**Figure 3:**
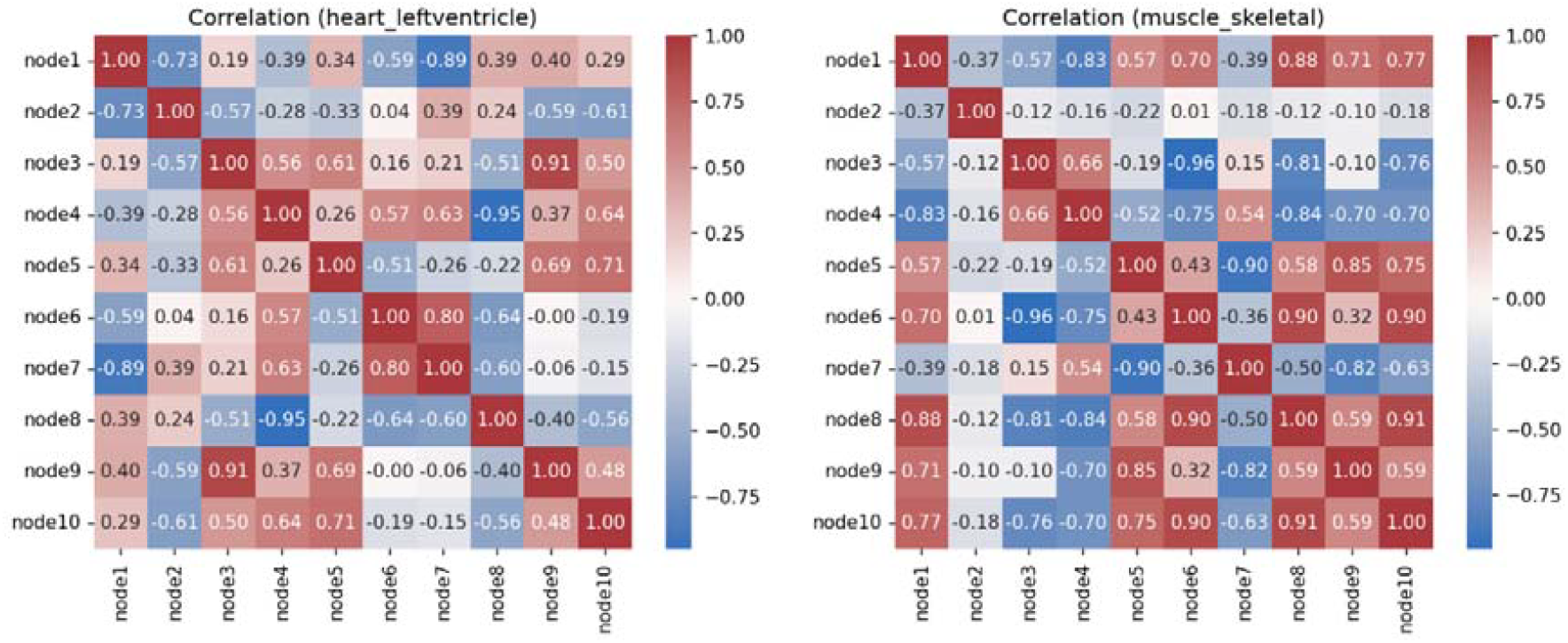
Tissue-specific latent node correlation matrices. (A) Heart – Left Ventricle (Group A) showing a mixture of strong positive and negative node–node associations. (B) Muscle – Skeletal (Group B) displaying more numerous high-magnitude correlations, including both strong positive couplings and pronounced antagonistic pairs.

Using decoder perturbations (+1.0 per latent dimension; core = Top-250 ∪ Top-5% by absolute reconstruction change), each node yielded a 250-gene core. Across ten nodes the union comprised 1,725 unique symbols, with modest pairwise overlap (mean intersection ≈23.6 genes; mean Jaccard ≈; 0.054), indicating largely non-redundant content, while the Jaccard similarity for node3–node4 was ≈0.43 (43%) with 152 shared genes, marking a notable exception. A small set of genes recurred across multiple cores: AZGP1 appeared in six cores (most frequent), and ALDOC, BEX3, DIRAS1, GPSM1, NDRG4, NEBL, PDE1C, SLC12A7, TAX1BP3 appeared in five cores each. Taken together, these recurrent genes point to shared programs typical of excitable/contractile tissues energy metabolism (glycolysis), G-protein/small-GTPase signaling, Ca^2+^-regulated cyclic-nucleotide turnover and ion homeostasis, and cytoskeletal/adhesion modules. Full per-node lists and pairwise overlaps are provided in the Supplementary Tables.

The enrichment profiles confirmed that the latent nodes captured coherent, tissue-relevant biology. Nodes 3–7 were strongly enriched for muscle contraction and cardiac conduction pathways, including striated muscle filament sliding and adrenergic signaling in cardiomyocytes. Other nodes mapped to complementary functions: Node 8 to mitochondrial energy metabolism and oxidative phosphorylation, Nodes 2 and 9 to extracellular matrix and adhesion, and Node 10 to ribosome biogenesis. These assignments indicate that the autoencoder disentangled the expression space into interpretable biological modules, rather than arbitrary feature sets. Importantly, disease-linked terms such as ventricular tachycardia, congenital myopathy, and arteriosclerosis emerged from relevant nodes, underscoring that the latent representations align with known pathophysiology.

Together, these results demonstrate that our approach not only separates tissues at the expression level but also recovers meaningful biological modules with direct functional and clinical relevance.

Applying QAOA to the tissue-specific latent correlation graphs yielded distinct outcomes for Heart and Muscle. For the Heart graph, 7 raw candidates were produced, but only 3 survived post-filtering. Each surviving subset was restricted to a simple 2-node pair with summed strong-edge weights just below 1.0 (0.95, 0.91, 0.89). Larger candidate subsets were eliminated during connectedness and deduplication checks. This fragmentation is reflected in the final Heart graph (Fig. 4A), where only three dyads persist, while most latent nodes remain isolated.

**Figure 4:**
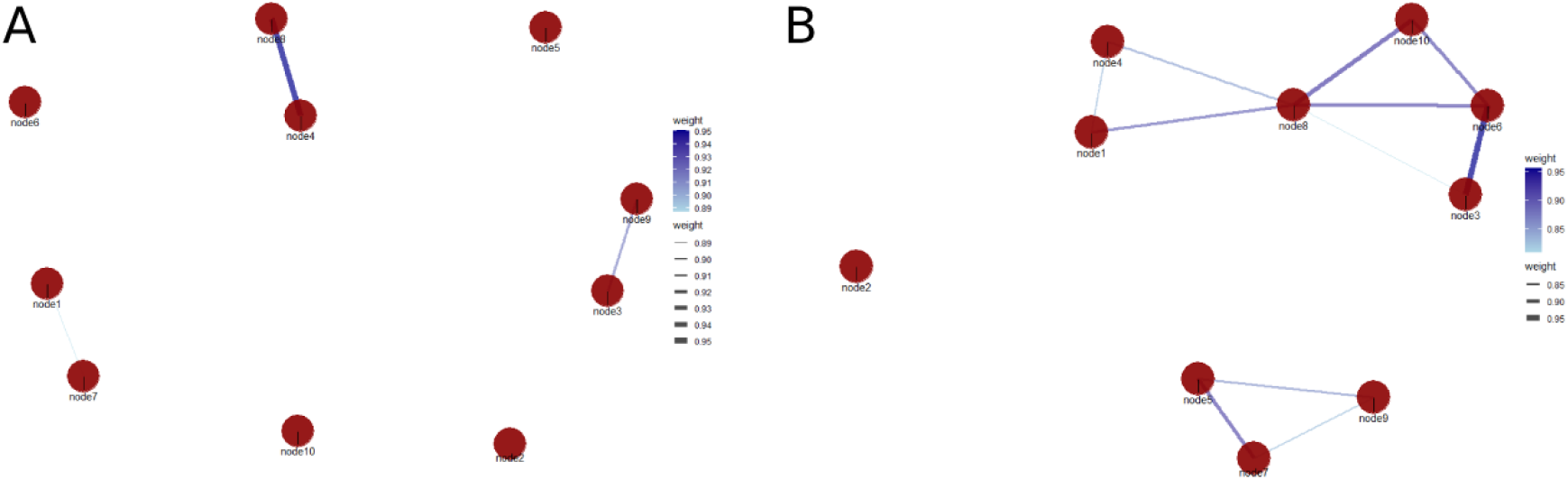
QAOA-derived modules from latent correlation graphs. (A) Heart – Left Ventricle: only three 2-node dyads (edges > 0.89) survived filtering, while most nodes remained isolated. (B) Muscle – Skeletal: one large, cohesive 6-node clique (nodes 1, 3, 4, 6, 8, 10) dominated the QAOA output. An additional 3-node grouping (nodes 5–7–9) appears visually but was excluded due to weaker edge weights (< top-10 threshold).

In contrast, the Muscle graph exhibited a markedly different organization. Out of 10 raw candidates, a single 6-node module passed all filters, supported by a dense web of strong edges (summed weight ≈ 7.0). This cohesive subgraph is shown in Fig. 4B, where nodes 1, 3, 4, 6, 8, and 10 form a tightly interconnected clique. Notably, an additional triplet (nodes 5–7–9) appears in the visualization but its edge weights fall below those recovered in the QAOA top-10 list. As a result, this weaker module was discarded, leaving the single dominant clique as the sole accepted subset.

Together, these results illustrate a clear tissue-specific contrast: the Heart latent graph fragments into multiple minimal dyads, whereas the Muscle latent graph converges into one large, dense, and biologically interpretable module. This inversion between sample-level heterogeneity (broader Muscle dispersion in latent PCA) and module-level cohesion (Muscle clique dominance) highlights the complementary nature of the autoencoder and quantum stages.

The QAOA-derived cliques align with coherent tissue programs. In Heart, the three surviving dyads (Fig. 4A) pair (i) conduction with mitochondrial metabolism (node4– node8), (ii) conduction with ECM/adhesion (node3–node9), and (iii) two contractile nodes (node1–node7). This pattern suggests tightly regulated pairwise coordination between excitation–contraction coupling, energy provision, and matrix context, rather than a single, extended module at the imposed edge threshold. In Muscle, the dominant 6-node clique (nodes 1, 3, 4, 6, 8, 10; Fig. 4B) integrates contractile machinery (nodes 1, 6), electrophysiology/conduction (nodes 3, 4), mitochondrial oxidative metabolism (node 8), and translation capacity (node 10), consistent with the high energetic and proteostatic demands of skeletal muscle. A secondary triangle (nodes 5–7–9) visible in Fig. 4B reflects a structural/ECM-leaning grouping; however, its edges are weaker and therefore it did not appear among the QAOA top-10 results, leaving the single integrated module as the Muscle consensus. Collectively, these cliques indicate modular dyadic coupling in Heart versus multicomponent integration in Muscle.

A distinctive feature of our formulation is that it does not enforce maximal cliques or strictly dense subgraphs, but instead retains any subset that is both connected and internally supported by strong edges. This subtle change is biologically important: it allows the solver to highlight connector nodes that bridge otherwise separate clusters. For example, in the Muscle graph, node 8 emerged as a linking element between two otherwise distinct groups, effectively acting as a bottleneck in the latent network. Such connector hubs are well known to mediate signal flow and regulatory cross-talk in real biology, yet they are often overlooked by clique-only or density-maximising heuristics. By combining a density reward with a non-isolation penalty, our pipeline recovers modules that better mirror the heterogeneous topologies observed in molecular systems, rather than restricting discovery to tightly packed cliques.

## Conclusion

We set out to test whether quantum optimization can add value to modern network biology. Our workflow first learns a 10-dimensional latent graph from expression data with an autoencoder, then uses QAOA to extract connected, high-weight modules. The results are clear and consistent with biology: in Heart we recovered three focused dyads (pairwise links between conduction, contractile, metabolism, and ECM programs), while in Muscle we found one integrated 6-node module combining contractile, electrophysiology, mitochondrial, and translational functions. This shows that quantum optimization can turn latent correlations into discrete, testable modules rather than diffuse patterns our intended goal.

Our study is a proof-of-concept and it worked under strict limits (10 nodes), but the method is not tied to this size. The same objective and post-filters can be scaled by building latent graphs hierarchically (coarse → fine), (ii) decomposing large graphs into blocks with overlap checks, and (iii) using hybrid classical–quantum warm starts. With these steps, the exact same idea can be pushed to thousands to hundreds of thousands of nodes as hardware and solvers improve.

The approach is also modality-agnostic. Edges in the graph can come from multi-omics (transcriptome, proteome, chromatin, metabolome) and from cross-layer links (e.g., TF–chromatin, protein–metabolite). In this way, quantum optimization becomes a general module finder for integrated biology, not just for a single data type.

To the best of our knowledge, this is one of the first attempts to apply a QAOA-style solver directly to learned biological latent graphs and to check the output against pathway-level meaning. Within this scope, we achieved our two main goals: (1) a clean, reproducible pipeline from data → latent graph → quantum modules, and (2) biologically coherent cliques that reflect known tissue programs. We therefore see quantum modeling as a credible option for the next generation of network biology promising today at small scale, and positioned to become transformative as scale and multi-omics integration grow.

## Acknowledgements

This work received no external funding. The authors thank all colleagues who provided feedback during manuscript preparation.

## Author contributions

- **Milad Yazdi:** Conceptualization; Methodology; Writing – Original Draft; Supervision; Writing – Review & Editing.
- **Kushala Ralpati Srinivas:** Data Curation; Software; Investigation; Validation; Writing – Review & Editing.
- **Ajay Yadav:** Methodology (mathematical formulation); Formal Analysis; Validation; Writing – Review & Editing.

## Data availability

The datasets analyzed in this study were obtained from the GTEx portal (https://gtexportal.org/). Processed results and code are available from the corresponding author upon reasonable request.

## Declarations

### Use of AI/LLMs

**Use of AI/LLMs** The authors used AI tools **only for language polishing**; all scientific content was written, reviewed, and approved by the authors, who take full responsibility for the article. **No AI-generated images** were used.

### Ethical approval

Not applicable.

### Informed consent

Not applicable.

